# Genome sequencing and functional genes comparison between *Sphingopyxis* USTB-05 and *Sphingomonas morindae* NBD5

**DOI:** 10.1101/2021.03.29.437629

**Authors:** Chao Liu, Qianqian Xu, Zhenzhen Zhao, Shahbaz Ahmad, Haiyang Zhang, Yufan Zhang, Yu Pang, Abudumukeyiti Aikemu, Yang Liu, Hai Yan

**Affiliations:** School of Chemistry and Biological Engineering, University of Science and Technology Beijing, Beijing 100083, China

**Keywords:** Sphingomonadaceae, Genome, Lutein, Hepatotoxin, Microcystins, Biodegradation

## Abstract

Sphingomonadaceae has a large number of strains that can biodegrade hepatotoxins or environmental pollutants. The latest research reported that certain strains can also produce lutein. Based on the third-generation sequencing technology, we analyzed the whole genome sequence and compared related functional genes of two strains of Sphingomonadaceae isolated from different habitats. The genome of *Sphingopyxis* USTB-05 was 4,679,489 bp and contained 4312 protein coding genes. The 4,239,716 bp nuclear genome of *Sphingomonas morindae* NBD5, harboring 3882 protein coding genes, has two sets of chromosomes. Both strains had lutein synthesis metabolism pathway sharing identical synthetic genes of *crtB*, *crtE*, *crtI*, *crtQ*, *crtL*, *crtR*, *atoB*, *dxs*, *dxr*, *ispD*, *ispE*, *ispDF*, *gcpE*, *ispG*, *ispH*, *ispA*, *ispB* and *ispU*. *Sphingopyxis* USTB-05 had hepatotoxins microcystins and nodularin metabolic pathways related to 16 genes (*ald*, *ansA*, *gdhA*, *crnA*, *phy*, *ocd*, *hypdh*, *spuC*, *nspC*, *speE*, *murI*, *murD*, *murC*, *hmgL*, *bioA* and *glsA*), while these genes were not found in *Sphingomonas morindae* NBD5. The unique protein sequences of strain NBD5 and strain USTB-05 were 155 and 199, respectively. The analysis of whole genome of the two Sphingomonadaceae strains provides insights into prokaryote evolution, the new pathway for lutein production and the new genes for environmental pollutant biodegradation.

**IMPORTANCE:** Understanding the functional genes related to the special functions of strains is essential for humans to utilize microbial resources. The ability of *Sphingopyxis* USTB-05 to degrade hepatotoxins microcystins and nodularin has been studied in depth, however the complete metabolic process still needs further elucidation. *Sphingomonas morindae* NBD5 can produce lutein, and it is necessary to determine whether there is a new pathway of lutein. In this study, the whole genome sequencing of *Sphingopyxis* USTB-05 and *Sphingomonas morindae* NBD5 were performed for the first time. Lutein synthesis metabolic pathways and synthetic genes were discovered in Sphingomonadaceae. We predicted the existence of new lutein synthesis pathways and revealed most of the genes of the new synthesis pathways. A comparative analysis of the functional genes of the two strains revealed that *Sphingopyxis* USTB-05 contains a large number of functional genes related to the biodegradation of hepatotoxins or hexachlorocyclohexane. Among them, the functional genes related to the biodegradation and metabolism of hexachlorocyclohexane had not been previously reported. These findings lay the foundation for the biosynthesis of lutein using *Sphingomonas morindae* NBD5 or *Sphingopyxis* USTB-05 and the application of *Sphingopyxis* USTB-05 for the biodegradation of hepatotoxins microcystins and nodularin or environmental pollutants.

## INTRODUCTION

Lutein is a kind of carotenoid, which widely exits in vegetables, fruits and other plant. It is also the main pigment in the macular area of human’s eyes (1), and cannot be synthesized by the body itself. Although it must be obtained from daily food, most people’s daily intake is seriously insufficient. The latest research showed that daily intake of 10 mg lutein and 2 mg zeaxanthin could improve visual function and delay the development of age-related macular degeneration (AMD) (2). Many researches focused on the production of lutein by eukaryotes, and its synthesis pathway had been clarified already. However, the production of lutein by prokaryotes was reported rarely. The prokaryotic strains were only confined to some specific strains, for examples, *Erwinia*, *Agrobacterium* and *Rhodobacter capsulatus* (2). Few research was reported about *Sphingomonas* strains in lutein production. In our previous research, *Sphingomonas morindae* NBD5 had the function of producing lutein (3). Phylogenetic analysis showed that they belonged to two closely related genera of Sphingomonadaceae (4) (5).

Sphingomonadaceae has a large number of strains that can biodegrade hepatotoxins or environmental pollutants. In the 1990s, *Sphingomonas wittichii* RW1 was reported to biodegrade dibenzo-p-dioxin (DD) polychlorinated derivatives under aerobic condition in contaminated soil and water (6). Other types of *Sphingomonas* could also biodegrade a series of intractable compounds (biphenyl, herbicide dichlorohaloperidin, γ-hexachlorocyclohexane, aromatic hydrocarbons, chlorophenol, pentachlorophenol, naphthalenesulfonic acid, N,N-dimethylaniline, diphenyl ether, dibenzofuran) that were harmful to the environment (7). Hepatotoxins microcystins (MCs) and nodularin (NOD) are derived from algae and have high toxicity and potential harm to humans and aquatic animals. The World Health Organization (WHO) stipulated that the concentration of MCs in drinking water should not be higher than 1.0 μg/L (8). Biodegradation is a very promising method to remove hepatotoxins. At present, many strains of Sphingomonadaceae family have the function of biodegrading hepatotoxins. However, in order to fully clarify the metabolic process of biodegradation, new hepatotoxins biodegradation genes need to be discovered.

*Sphingomonas morindae* NBD5 was a new species that was identified as this genus only a few years ago (4). It could produce high yield of lutein, and the metabolic genes needed further study. *Sphingopyxis* USTB-05 was isolated from Dianchi Lake in China and could biodegrade MCs and NOD (5) (9) (10). These functional genes *USTB-05-A*, *USTB-05-B*, and *USTB-05-C* of *Sphingopyxis* sp. USTB-05 had been verified by heterologous expression in *Escherichia coli* to biodegrade MCs (11)(12). The purified first recombinant enzyme was found to have a strong ability to catalyze hepatotoxins in *Sphingopyxis* sp. USTB-05 (13). Because of the characteristics of lutein production and hepatotoxin biodegradation of these two strains, the relationship between gene and function was found through searching for functional genes. Here, the method of whole genome combined with biometric analysis was used to compare the similar and unique characteristics of *Sphingomonas morindae* NBD5 and *Sphingopyxis* USTB-05, and analyze their functional genes, especially those associated with lutein synthesis and hepatotoxin biodegradation.

## RESULTS

### General features of the nuclear genome

The *Sphingomonas morindae* NBD5 genome contained two circular chromosomes and two circular plasmids (Figure 1). Polychromosomes were common in some genera, but rare in *Sphingomonas*. Two circular chromosome of 4,239,716 bases was finally obtained with a G + C content of 70%, and 3882 protein coding sequences (CDSs) were predicted, accounting for 62.39% of the total coding sequence. The 16S rRNA of strain NBD5 had 3 complete copies (Table 1). The *Sphingopyxis* USTB-05 genome contained one chromosome (Figure 2). Its circular chromosome of 4,679,489 bases was finally obtained with a G + C content of 64%, and 4312 CDSs were predicted, accounting for 62.39% of the total coding sequence. The 16S rRNA of strain USTB-05 was a single copy (Table 1) without CRISPR site. Compared with strain USTB-05, the genome of strain NBD5 was much richer in GC and contained plasmids, which indicated that some genetic transfer events happened in strain NBD5 (Table 1). The number of functional genes annotated in strain NBD5 was much more than that in strain USTB-05, but the chromosome length of strain USTB-05 was longer than that of strain NBD5.

**Figure 1:**
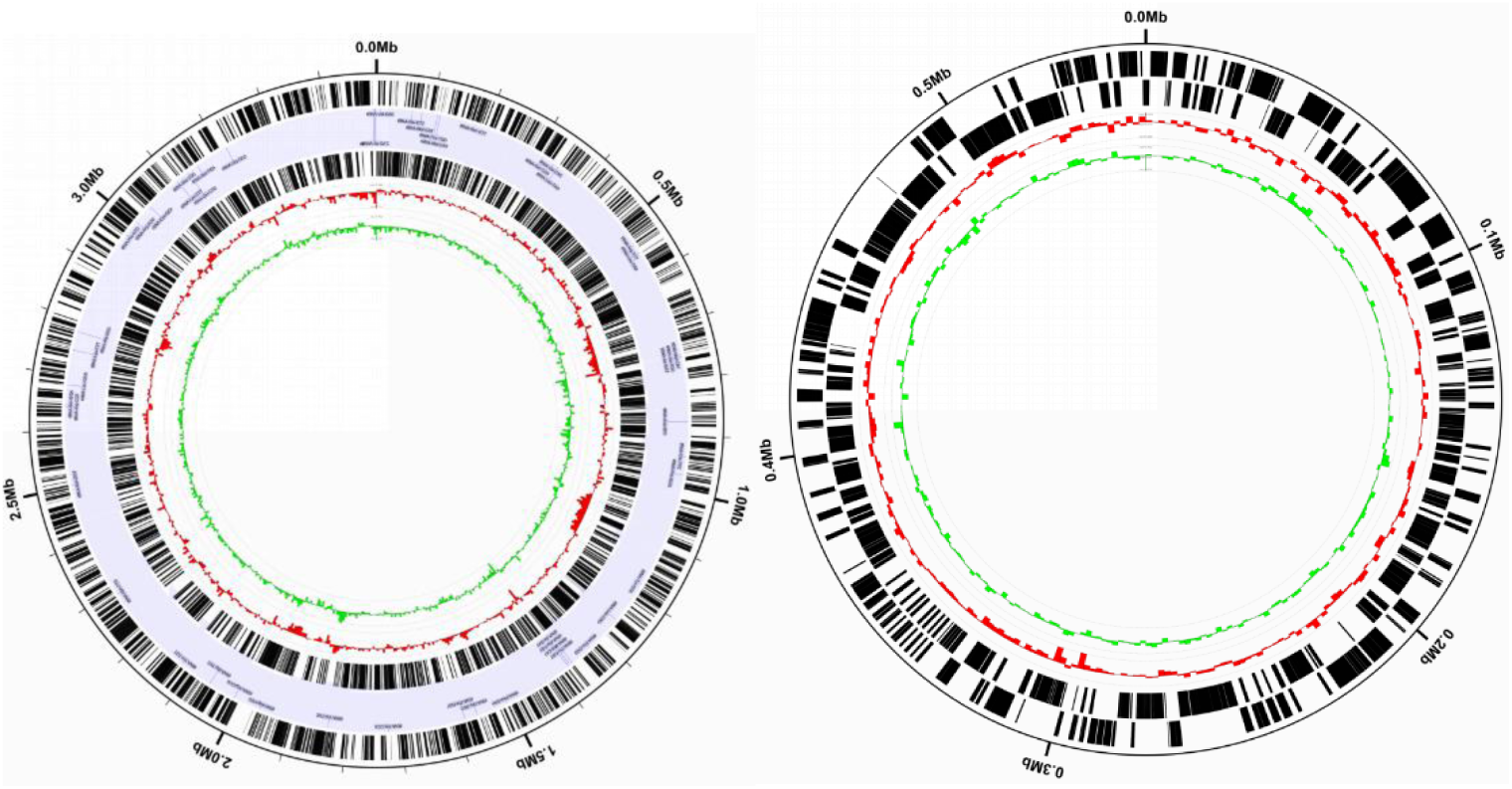
Genome circle of *Sphingomonas morindae* NBD5.

**Figure 2:**
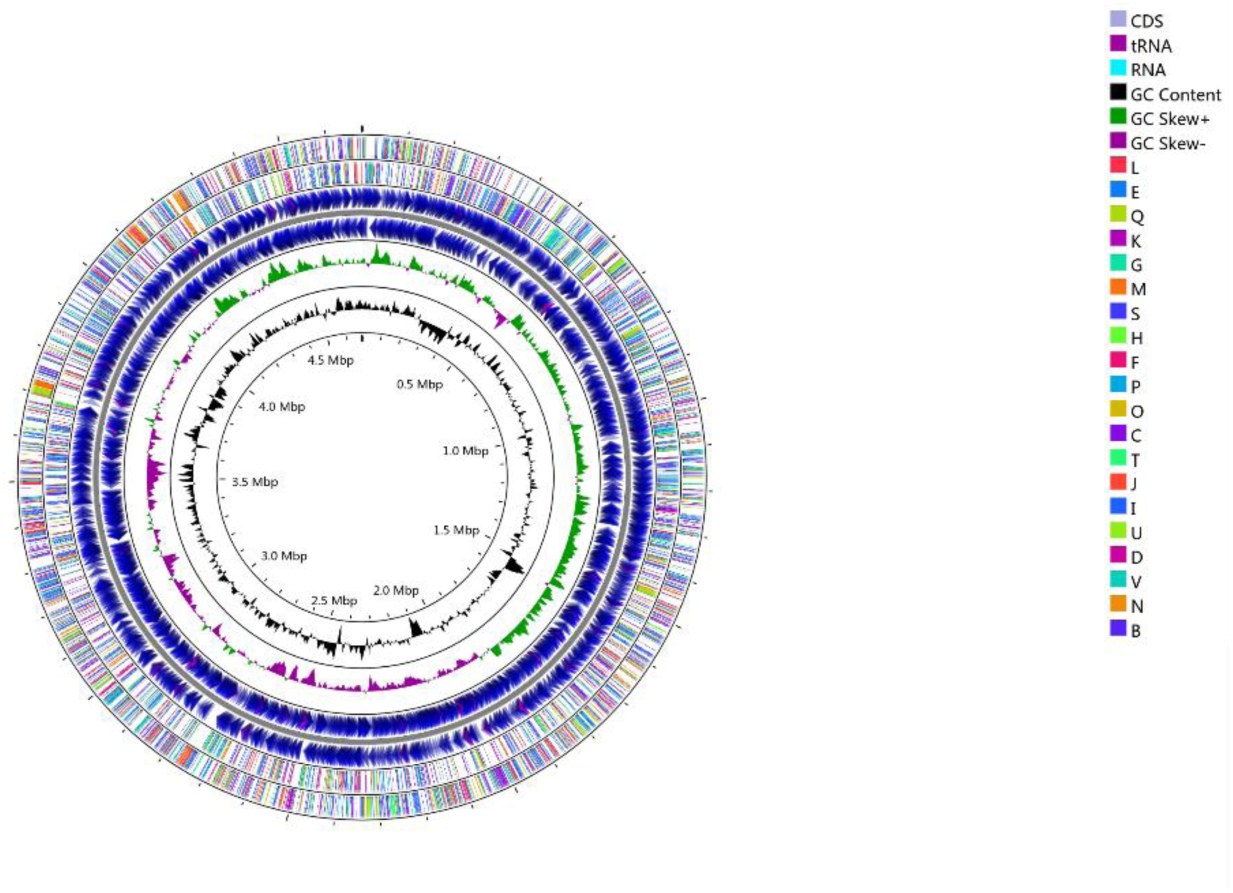
Genome circle of *Sphingopyxis* USTB-05.

**Table 1.** Comparison of genome characteristics between *Sphingomonas morindae* NBD5 and *Sphingopyxis* USTB-05.

| category | *Sphingomonas morindae* NBD5 | *Sphingopyxis* USTB-05 |
| --- | --- | --- |
| bases | 4239716 | 4679489 |
| tmRNA | 1 | 1 |
| tRNA | 61 | 48 |
| CDS | 3882 | 4312 |
| GC(%) | both 70% | 64% |
| plasmid | 2 | 0 |

### Genome annotation and genome wide comparative analysis

#### GO annotation

The distribution of genes in different Gene Ontology (GO) terms can intuitively reflect the distribution of target genes on the secondary level of GO terms. A total of 798 genes were annotated in the *Sphingomonas morindae* NBD5 genome through the GO database (Figure 3). Under the classification of biological process, the number of genes annotated to metabolic process was up to 43.8%, and the number of genes for intracellular processes was 40.8%. Response to stimulus, biological regulation, localization, regulation of biological process, multi-organism process, cellular component organization or biogenesis, signaling, and growth accounted for 15%, 14%, 11.8%, 11.8%, 7.3%, 6.3%, 4.1%, and 3%, respectively. Under the classification of cellular component, the number of genes annotated to cell was up to 38%, and the number of genes for cell part was 37.9%. Membrane, membrane part, protein-containing complex, organelle, extracellular region, organelle part, nucleoid, and other organism accounted for 22.6%, 14.8%, 6.2%, 3.2%, 1.9%, 1.5%, 0.4%, and 0.2%, respectively. Under the classification of molecular function, the number of genes annotated to catalytic activity was up to 1623 (40.6%), and the number of genes for binding was 37.5%. Transporter activity, transcription regulator activity, structural molecule activity, molecular transducer activity, obsolete signal transducer activity, antioxidant activity, molecular function regulator, and molecular carrier activity accounted for 6.9%, 4.1%, 1.7%, 1.6%, 1.4%, 0.7%, 0.4%, and 0.1%, respectively.

**Figure 3:**
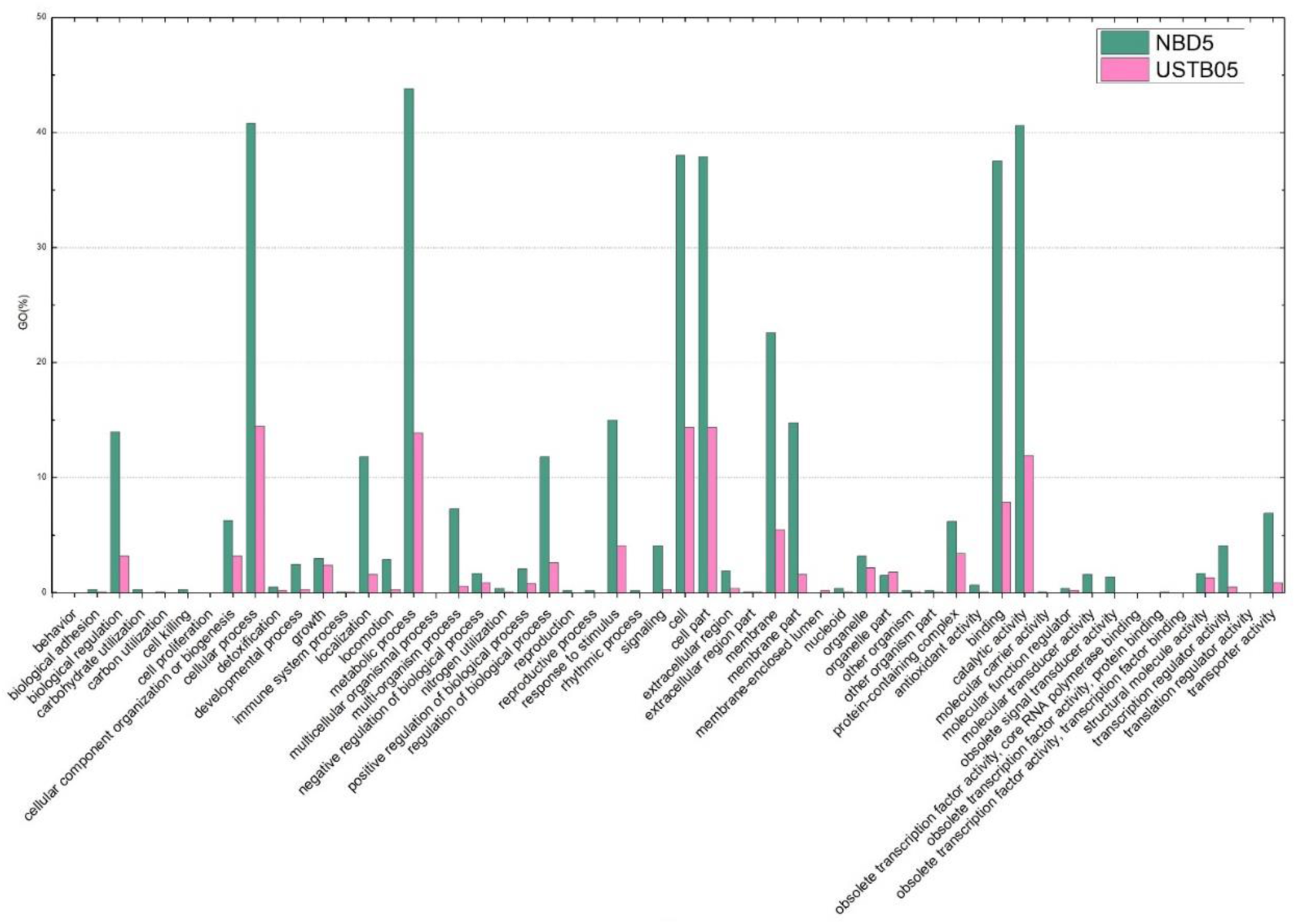
Comparison of GO functional classification of *Sphingomonas morindae* NBD5 and *Sphingopyxis* USTB-05.

A total of 786 genes were annotated in the *Sphingopyxis* USTB-05 genome through the GO database (Figure 3). Under the classification of biological process, the number of genes annotated to cellular process was up to 14.5%, and the number of genes for metabolic process was 13.9%. Response to stimulus, cellular component organization or biogenesis, biological regulation, regulation of biological process, growth, localization, negative regulation of biological process, and positive regulation of biological process accounted for 4.1%, 3.2%, 3.2%, 2.6%, 2.4%, 1.6%, 0.9%, and 0.8%, respectively. Under the classification of cellular component, the number of genes annotated to cell was up to 14.4%, and the number of genes for cell part was 14.4%. Membrane, protein-containing complex, organelle, organelle part, membrane part, extracellular region, membrane-enclosed lumen, and extracellular region part accounted for 5.5%, 3.4%, 2.2%, 1.8%, 1.6%, 0.4%, 0.2%, and 0.1%, respectively.

Under the classification of molecular function, the number of genes annotated to catalytic activity was up to 11.9%, and the number of genes for binding was 7.9%. Structural molecule activity, transporter activity, transcription regulator activity, molecular function regulator, and antioxidant activity accounted for 1.3%, 0.9%, 0.5%, 0.2%, and 0.1%, respectively.

Although *Sphingopyxis* USTB-05 had more bases in its genome than *Sphingomonas morindae* NBD5, *Sphingomonas morindae* NBD5 had significantly more genes in most GO classifications than *Sphingopyxis* USTB-05 (Figure 3). *Sphingomonas morindae* NBD5 had 66 unique genes in carbohydrate utilization (GO:0009758): 11 and obsolete signal transducer activity (GO:0004871): 55. But there were also some genes in behavior (GO:0007610): 1, cell proliferation (GO:0008283): 2, multicellular organismal process (GO:0032501): 2, obsolete transcription factor activity, protein binding (GO:0000988): 4, obsolete transcription factor activity, transcription factor binding (GO: 0000989): 2, translation regulator activity (GO: 0045182): 1, these genes in *Sphingopyxis* USTB-05 were unique, representing behavior; cell proliferation; multicellular organismal process; obsolete transcription factor activity, protein binding; obsolete transcription factor activity, transcription factor binding; translation regulator activity, respectively.

### COG annotation

Cluster of Orthologous Groups (COG) is a database based on the systematic evolution of bacteria, algae and eukaryotes. The assembled *Sphingomonas morindae* NBD5 gene was analyzed in the COG database, and the results showed that 62.39% of the genes in COG were annotated (Figure 4). In the 25 COG functional categories, 3668 genes had been classified. These categories were mainly: transcription (COG category K) (7.66%, as a percentage of all functional allocation genes), cell wall/membrane/envelope biogenesis (M) (7.06%), carbohydrate transport and metabolism (G) (6.82 %), amino acid transport and metabolism (E) (6.73%), signal transduction mechanisms (T) (6.05%), inorganic ion transport and metabolism (P) (5.62%), energy production and conversion (C) (5.37%), translation, ribosomal structure and biogenesis (J) (4.80%), replication, recombination and repair (L) (4.77%), and lipid transport and metabolism (I) (4.58%). The rate of COG classification as transcription (K) was high (7.66%), which was consistent with the fact that this strain contained two circular chromosomes and complex gene expression.

**Figure 4:**
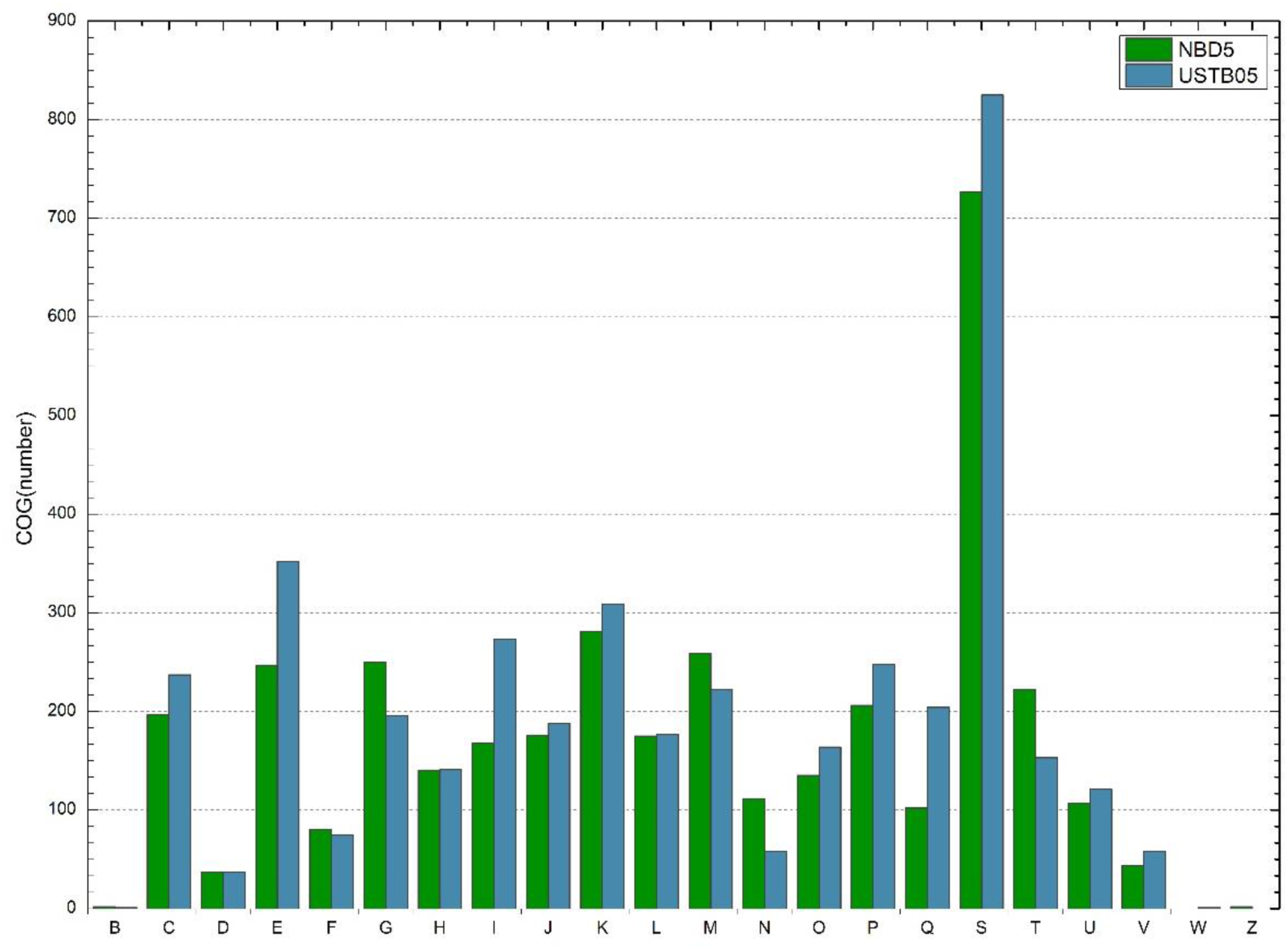
Comparison of COG functional classification of *Sphingomonas morindae* NBD5 and *Sphingopyxis* USTB-05.

Analysis of the assembled *Sphingopyxis* USTB-05 gene in the COG database showed that 62.39% of the genes were annotated in COG. Among the 25 functional categories of COG, a total of 4,040 genes had been classified (Figure 4). These categories were mainly: amino acid transport and metabolism (E) (8.71%), transcription (K) (7.65%), lipid transport and metabolism (I) (6.76%), inorganic ion transport and metabolism (P) (6.14%), energy production and conversion (C) (5.87%), cell wall/membrane/envelope biogenesis (M) (5.50%), secondary metabolite biosynthesis, transport and catabolism (Q) (5.05%), carbohydrate transport and metabolism (G) (4.85%), translation, ribosomal structure and biogenesis (J) (4.65%), replication, recombination and repair (L) (4.38%), and posttranslational modification, protein turnover, chaperones (O) (4.06%).

Most COGs showed similar distributions among *Sphingomonas morindae* NBD5 and *Sphingopyxis* USTB-05 (Figure 4). Among them, *Sphingopyxis* USTB-05 had the largest number of genes in most COGs categories; however, the number of genes in the COG G, M, and T categories for the *Sphingopyxis* USTB-05 was lower. The COG G, M, and T categories for *Sphingomonas morindae* NBD5 represented carbohydrate transport and metabolism, cell wall/membrane/envelope biogenesis and signal transduction mechanisms, respectively. These data suggested that *Sphingomonas morindae* NBD5 had significant differences in carbohydrate-related metabolic synthesis and signal transduction. These reflected the fact that *Sphingomonas morindae* NBD5 could produce more lutein than *Sphingopyxis* USTB-05, and the synthesis of lutein was related to the metabolic synthesis of carbohydrates. The number of genes in the COG C, E, I, and Q categories for *Sphingopyxis* USTB-05 was higher; they represented energy production and conversion; amino acid transport and metabolism; lipid transport and metabolism and secondary metabolites biosynthesis, transport and catabolism, respectively. These data suggested that *Sphingopyxis* USTB-05 had significant differences in catabolism. These reflected the fact that *Sphingopyxis* USTB-05 had the molecular basis for decomposing MCs that a cyclic polypeptide composed of seven amino acids in the environment.

#### KEGG annotation

Biological functions usually require coordination of different genes. Therefore, in order to identify the representative biological pathway of strain NBD5, single genes were annotated. A total of 1,839 genes were annotated as 112 metabolic pathways, some of which could be matched with multiple metabolic pathways. In the Kyoto Encyclopedia of Genes and Genomes (KEGG) database, genes annotated as 23 major pathways accounted for more than half of all annotated genes. The top ten pathways were: amino acid biosynthesis, carbon metabolism, two-component system, purine metabolism, flagella assembly, oxidative phosphorylation, ribosomes, ABC transporter, bacterial chemotaxis, and pyrimidine metabolism (Figure 5). These annotations provided important information about the specific biological processes and pathways of strain NBD5.

**Figure 5:**
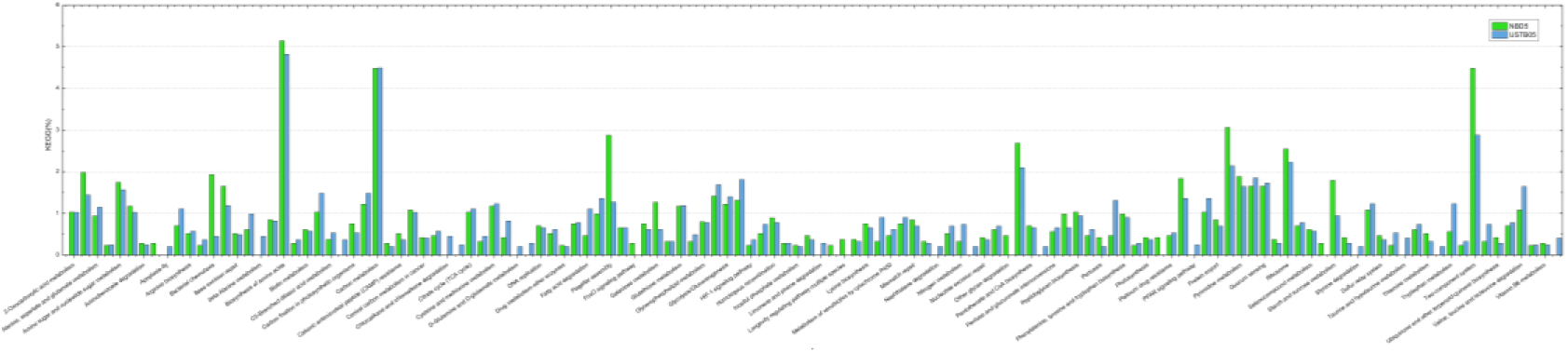
Comparison of KEGG pathway distribution of *Sphingomonas morindae* NBD5 and *Sphingopyxis* USTB-05.

A total of 1927 genes of strain USTB-05 had been annotated to 121 metabolic pathways, some of which could be matched with multiple metabolic pathways. In the KEGG database, genes annotated to 28 major pathways accounted for more than half of all annotated genes. The top ten pathways were: amino acid biosynthesis, carbon metabolism, two-component system, ribosomes, purine metabolism, oxidative phosphorylation, Pyruvate metabolism, glyoxylate and dibasic acid metabolism, quorum sensing, glycine, and serine and threonine metabolism (Figure 5). These notes provided important information about the specific biological processes and pathways of strain USTB-05.

Comparative analysis of KEGG function of the two strains of Sphingomonadaceae was did as well. Among the top ten pathways, two-component system (ko02020), purine metabolism (ko00230), flagella assembly (ko02040) (correlation of the neatness of colony edges on the plate), bacterial chemotaxis (ko02030)(correlation of the neatness of colony edges on the plate) genes strain NBD5 was obviously more than strain USTB-05; and amino acid biosynthesis (ko01230)(synthesis of enzymes related to algal toxin degradation), carbon metabolism (ko01200)(synthesis of enzymes related to algal toxin degradation), pyruvate metabolism (ko00620), glyoxylic acid and dibasic acid metabolism (ko00630), quorum sensing (ko02024), glycine, serine and threonine metabolism (ko00260)(synthesis of enzymes related to the degradation of algal toxins). The process gene strain USTB-05 was obviously more than strain NBD5. In addition, the KEGG function annotated that the unique metabolic processes of strain NBD5 included other polysaccharide degradation (ko00511), plant-pathogen interaction (ko04626) (from endophytes in plant noni), life regulation pathways-multi-species (ko04213), FoxO signaling pathway (ko04068), sphingolipid metabolism (ko00600), life regulation pathway (ko04211); the unique metabolic processes of strain USTB-05 included β-alanine metabolism (ko00410), degradation of chlorinated alkanes and chloroalkenes (ko00625), taurine and hypotaurine metabolism (ko04626), xylene degradation (ko00622), caprolactam degradation (ko00930), dioxin degradation (ko00621), limonene and pinene degradation (ko00903), chlorocyclohexane and chlorobenzene degradation (ko00361), PPAR signaling pathway (ko03320), apoptosis (ko04214), D-glutamine and D-glutamate metabolism (ko00471), naphthalene degradation (ko00626), styrene degradation (ko00643), toluene degradation (ko00623). This was consistent with the fact that the strain NBD5 was an endophyte and lutein-producing functional bacteria from the plant Noni, while the strain USTB-05 was a functional bacteria that could biodegrade complex organic matter in the environment.

### Comparison of the biosynthesis pathways and genes of lutein

Lutein contains two ionone rings in its chemical formula and is a carotenoid with vitamin A activity. *Sphingomonas morindae* NBD5 genome COG analysis shows that its core carbon skeleton is completed by carbohydrate transport and metabolism (G) (6.82%). Among them, the ratio of cell wall/membrane/envelope biogenesis (M) (7.06%) related to compound endocytosis and exocytosis is significantly higher. In addition, energy production and conversion (C) (5.37%) also plays an important role (Figure 4).

The corresponding genes of the terpenoid backbone biosynthesis pathway and the carotenoid biosynthesis pathway of *Sphingomonas morindae* NBD5 and *Sphingopyxis* USTB-05 were the same. The corresponding enzyme genes for lutein synthesis were also the same. Both strains found the presence of β-carotene 3-hydroxylase, which was involved in the last step of lutein synthesis, and its corresponding code gene was *CrtR-b*. This is a fact that lycopene existed as a synthetic intermediate, which is synthesized through glycolysis/gluconeogenesis pathway (Figure 6), terpenoid backbone biosynthesis pathway (Figure 7) and carotenoid biosynthesis pathway (Figure 8). But in the metabolic pathway of lutein production, no complete synthesis pathway had been found. Combined with the mass spectrometry results (3), the two strains of Sphingomonadaceae could synthesize lutein, which suggest that they could probably find new pathways of lutein synthesis.

**Figure 6:**
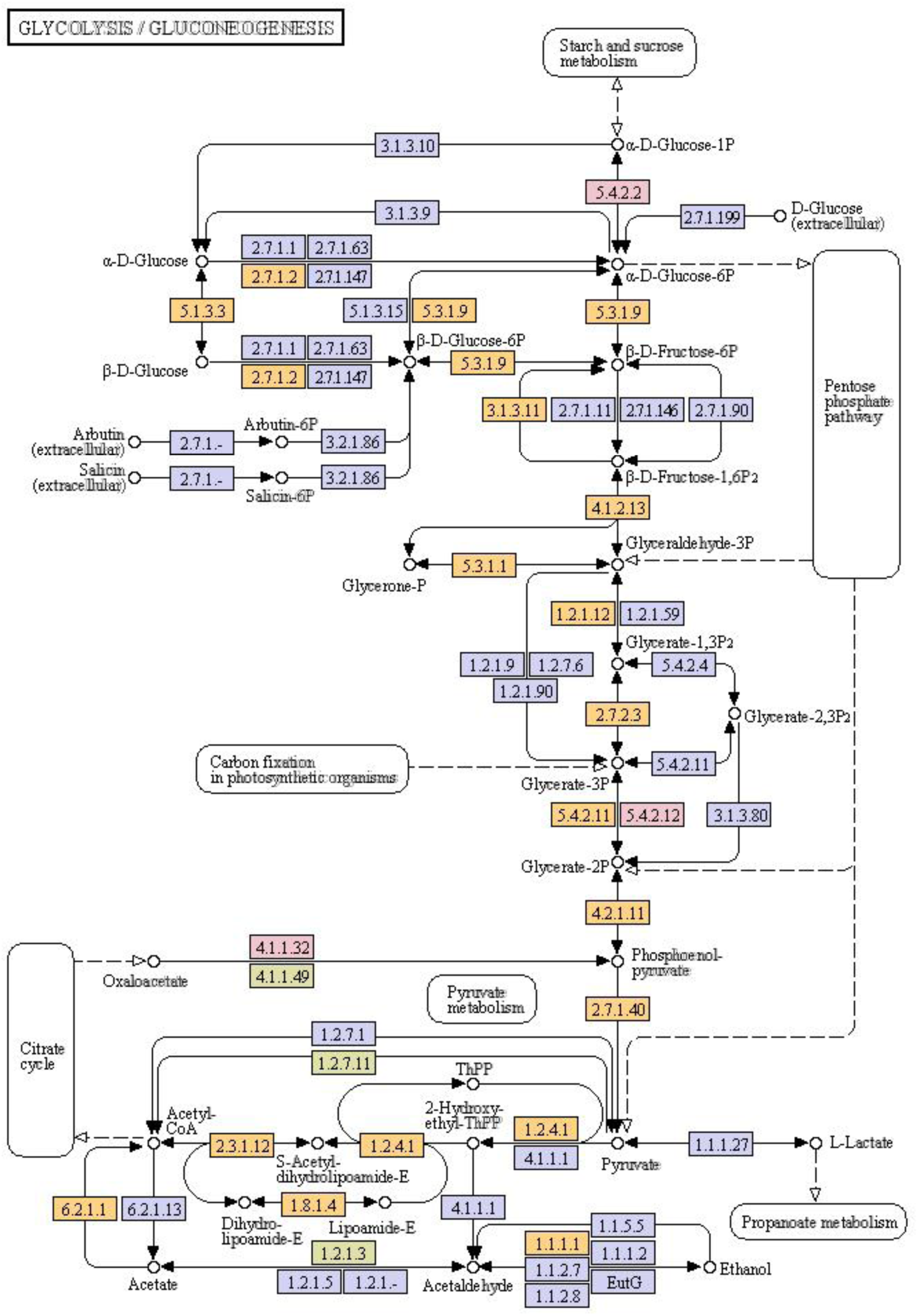
The metabolic pathway of glycolysis/gluconeogenesis of *Sphingomonas morindae* NBD5 and *Sphingopyxis* USTB-05. The enzyme or gene in the orange box in the figure indicates that two strains co-exist, the enzyme or gene in the blue box indicates that neither of the two bacteria exist, the enzyme or gene in the pink box indicates that it only exists in *Sphingomonas morindae* NBD5, and the enzyme or gene in the bright yellow box indicates that it only exists in *Sphingopyxis* USTB-05.

**Figure 7:**
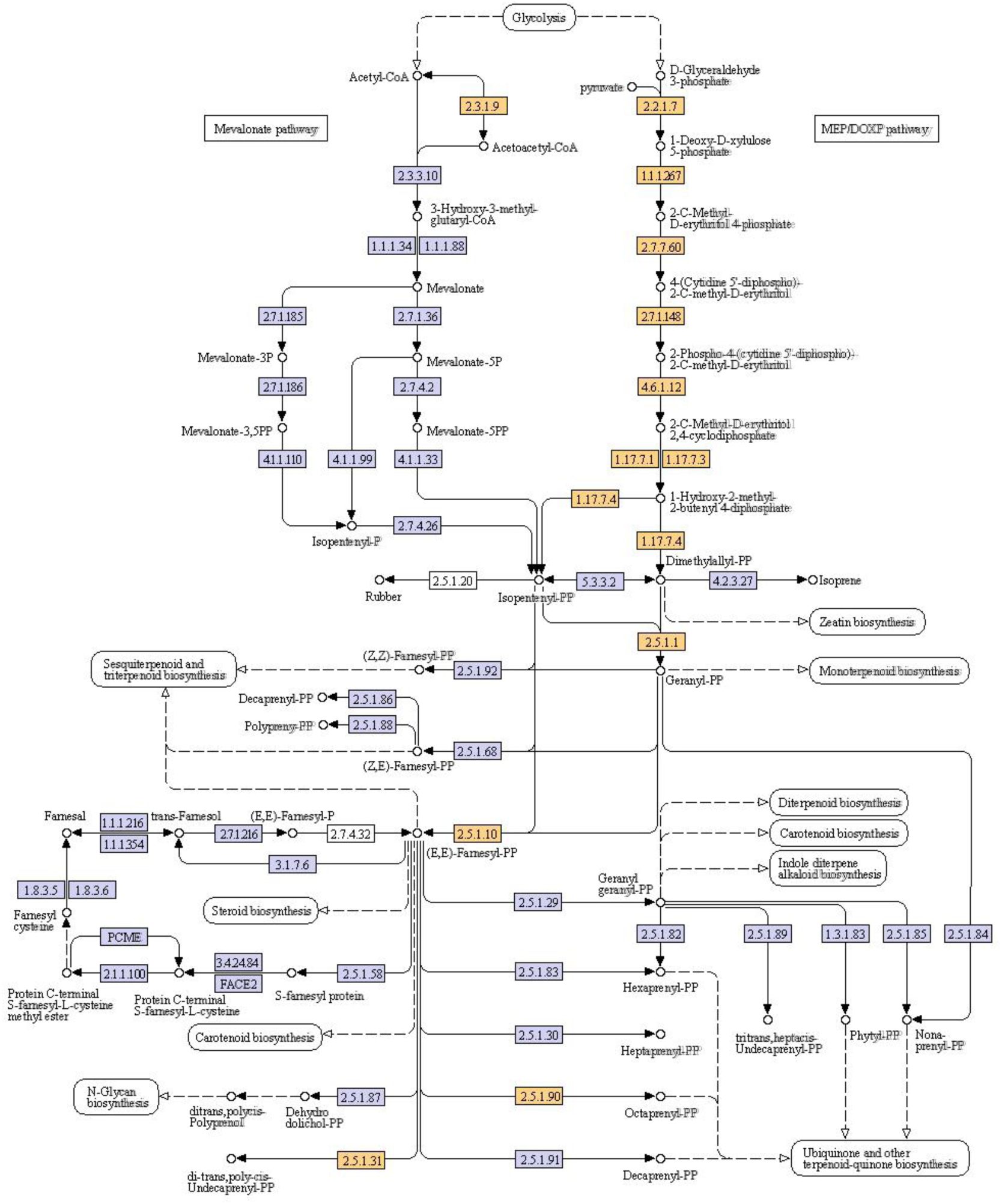
The biosynthetic pathway of terpenoid skeleton of *Sphingomonas morindae* NBD5 and *Sphingopyxis* USTB-05. The enzyme or gene in the orange box in the figure indicates that two strains co-exist, and the enzyme or gene in the blue box indicates that neither of the two bacteria exist.

**Figure 8:**
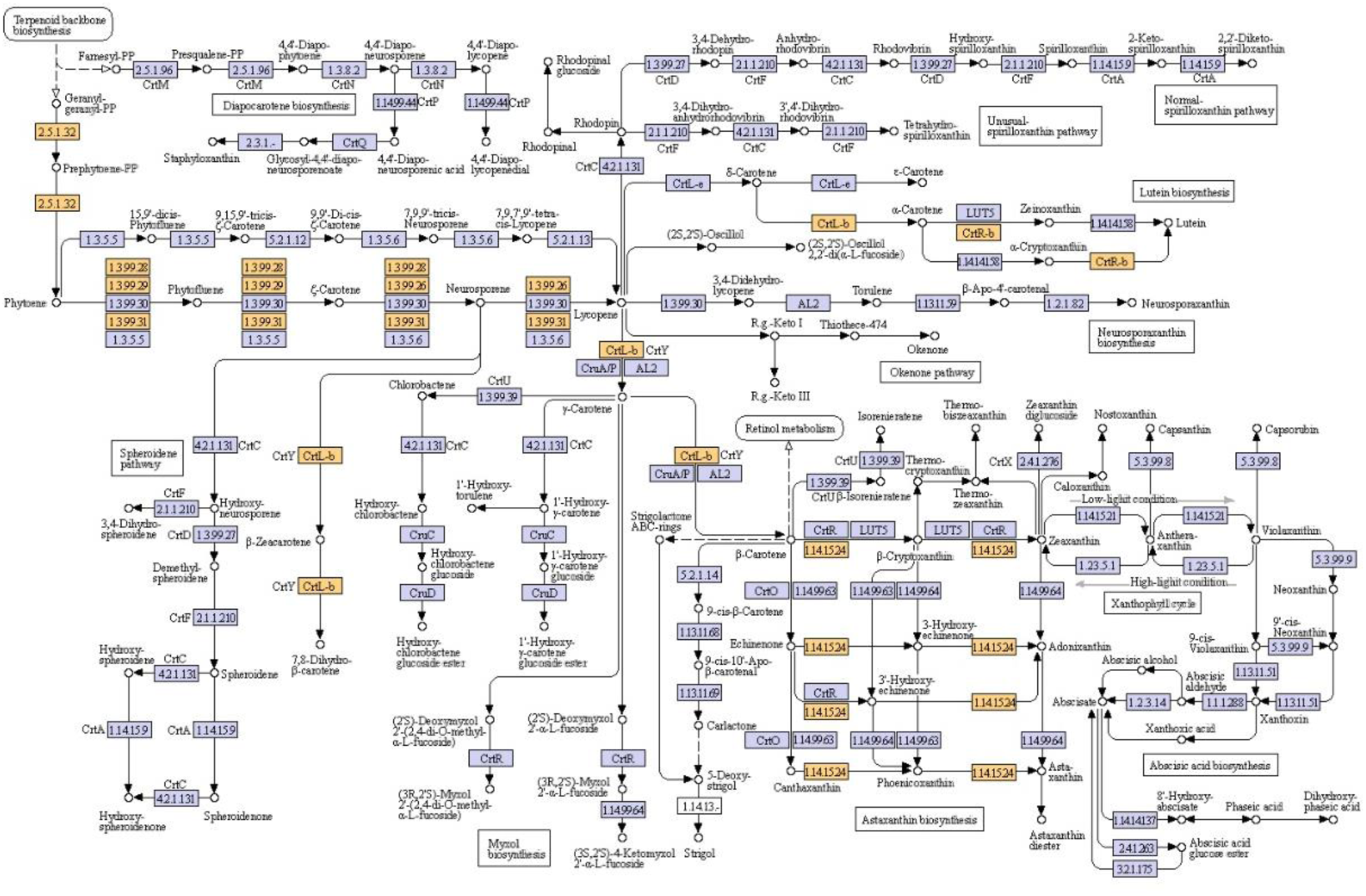
The biosynthetic pathway of carotenoid of *Sphingomonas morindae* NBD5 and *Sphingopyxis* USTB-05. The enzyme or gene in the orange box in the figure indicates that two strains co-exist, and the enzyme or gene in the blue box indicates that neither of the two bacteria exist.

In eukaryotes, especially higher plant, lutein is primarily synthesized from terpenoids. Two molecules of geranylgeranyl diphosphate (GGPP) are used to synthesize phytoene. Phytoene is then converted to ζ-carotene, after which ζ-carotene is converted into lycopene. Lycopene is then converted to α-carotene through cyclization reaction catalysed by lycopene cyclase (εLCY /βLCY). Finally, the formation of lutein from α-carotene is catalysed by α-carotene hydroxylase (βCHX /εCHX) (14). In the carotenoid biosynthesis pathway of the two Sphingomonadaceae strains, enzymes related to lycopene synthesis have been found (Figure 8), but the cyclases from lycopene to lutein synthesis are lacking. Therefore, it inferred that there were new cyclase genes for the lutein metabolism pathways in these two bacteria.

The lutein synthesis genes in *Sphingomonas morindae* NBD5 and *Sphingopyxis* USTB-05 genomes mainly existed in the terpenoid backbone biosynthesis pathway and the carotenoid biosynthesis pathway, which are completely the same. These synthetic genes are *crtB*, *crtE*, *crtI*, *crtQ*, *crtL*, *crtR*, *atoB*, *dxs*, *dxr*, *ispD*, *ispE*, *ispDF*, *gcpE*, *ispG*, *ispH*, *ispA*, *ispB* and *ispU*. Only these genes *ackA*, *pgm*, *gpm*I, and *pck*A in the glycolysis & gluconeogenesis pathway in *Sphingomonas morindae* NBD5 are unique. These genes *porB*, *meh*, and *fldA* in the glycolysis & gluconeogenesis pathway in *Sphingopyxis* USTB-05 are unique (Figure 9).

**Figure 9:**
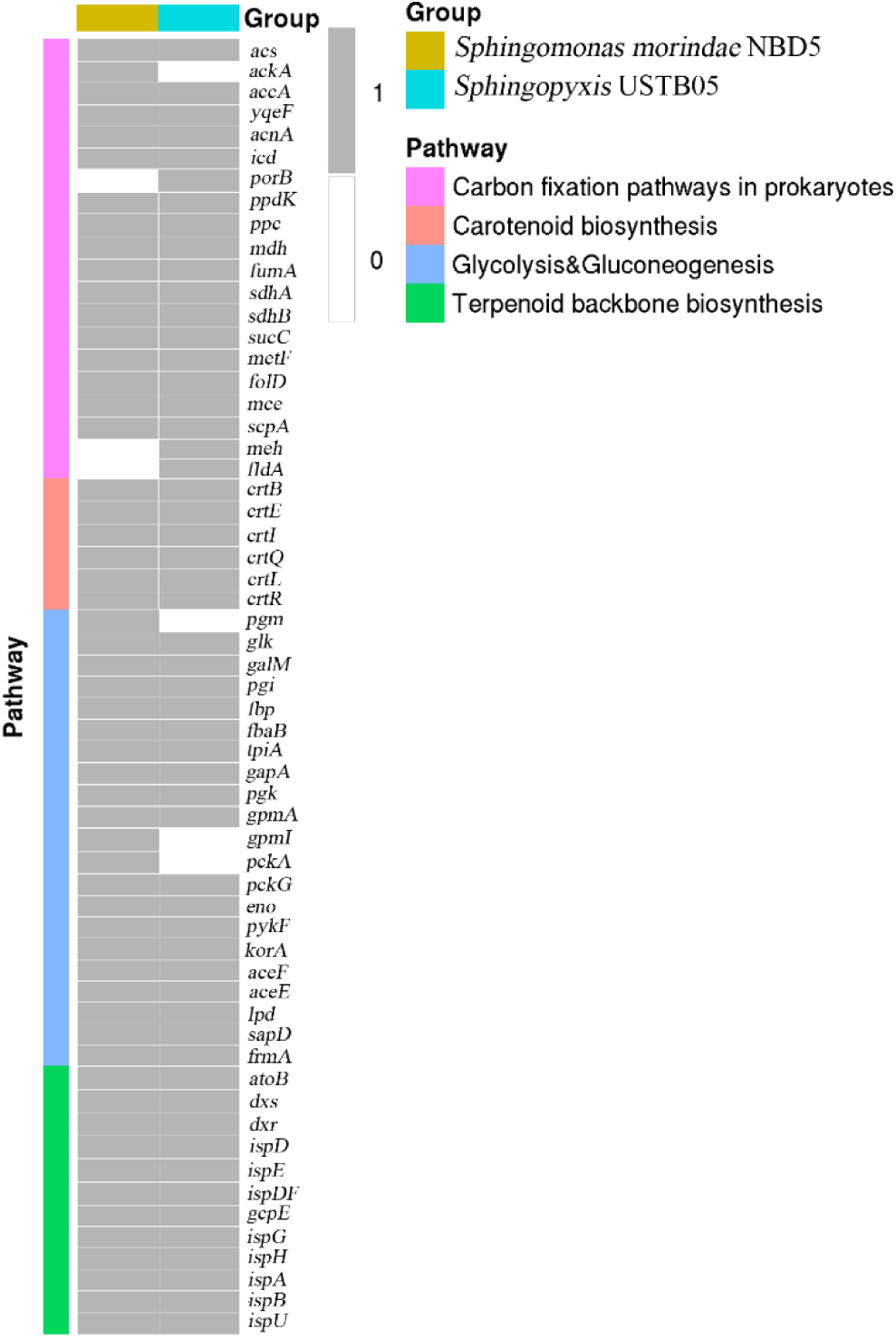
Comparison of genes related to lutein synthesis in the genomes of *Sphingomonas morindae* NBD5 and *Sphingopyxis* USTB-05. The gray box in the figure indicate that the gene exists in the strain, and the white indicate that the gene does not exist in the strain.

### Comparison of the pathways and genes of hepatotoxin biodegradation

Nowadays, the *mlr* gene cluster encoding functional hydrolase have been identified in many hepatotoxin biodegrading bacteria (15). The *mlr* gene cluster contained *mlrC*, *mlrA*, *mlrB* and *mlrD*, corresponding to the enzymes MlrC, MlrA, MlrB and MlrD that biodegraded hepatotoxin. By comparing the genomes of these two strains, it was found that *Sphingopyxis* USTB-05 contained the high homology genes *mlrC*, *mlrA*, *mlrB* and *mlrD*, while *Sphingomonas morindae* NBD5 did not have the *mlr* gene cluster (Suppl Table S1). It was consistent with the fact that *Sphingopyxis* USTB-05 was MCs biodegrading strain, while *Sphingomonas morindae* NBD5 was not.

Since hepatotoxin MCs and NOD are a class of monocyclic heptapeptide and pentapeptide compounds, some amino acid metabolism processes may be involved in MCs and NOD biodegradation. According to the general chemical molecular structure of MCs and NOD, these structures are D-alanine, variable L-amino acid, D-isoleucine, D-erythro-β-methylaspartic acid, N-dehydrogenation alanine, L-arginine, D-glutamic acid, Adda. Adda is the particular C20 β-amino acid: (2*S*, 3*S*, 8*S*, 9*S*) 3-amino-9-methoxy-2, 6, 8-trimethyl-10-phenyldeca-4(*E*), 6(*E*)-dienoic acid (10). Among them, the variable L-amino acids are leucine and arginine. *Sphingopyxis* USTB-05 had the following metabolic processes: alanine, aspartate and glutamate metabolism; arginine and proline metabolism; degradation of aromatic compounds; valine, leucine and isoleucine biosynthesis; D-glutamine and D-glutamate metabolism.

These MCs and NOD biodegradation genes in the genome of *Sphingopyxis* USTB-05 mainly existed in ABC transporters; alanine, aspartate and glutamate metabolism; arginine and proline metabolism; D-glutamine and D-glutamate metabolism; and valine, leucine and isoleucine degradation. These genes *ald*, *ansA*, and *gdhA* in the metabolic pathways of alanine, aspartate and glutamate were unique to *Sphingopyxis* USTB-05. These genes *crnA*, *phy*, *ocd*, *hypdh*, *spuC*, *nspC*, and *speE* in the metabolic pathways of arginine and proline were unique to *Sphingopyxis* USTB-05. These genes *murI*, *murD*, and *murC* in the metabolic pathways of D-glutamine and D-glutamate were unique to *Sphingopyxis* USTB-05. These genes *hmgL*, *bioA*, and *glsA* in the degradation pathway of valine, leucine and isoleucine may be involved in the biodegradation of MCs (Figure 10).

**Figure 10:**
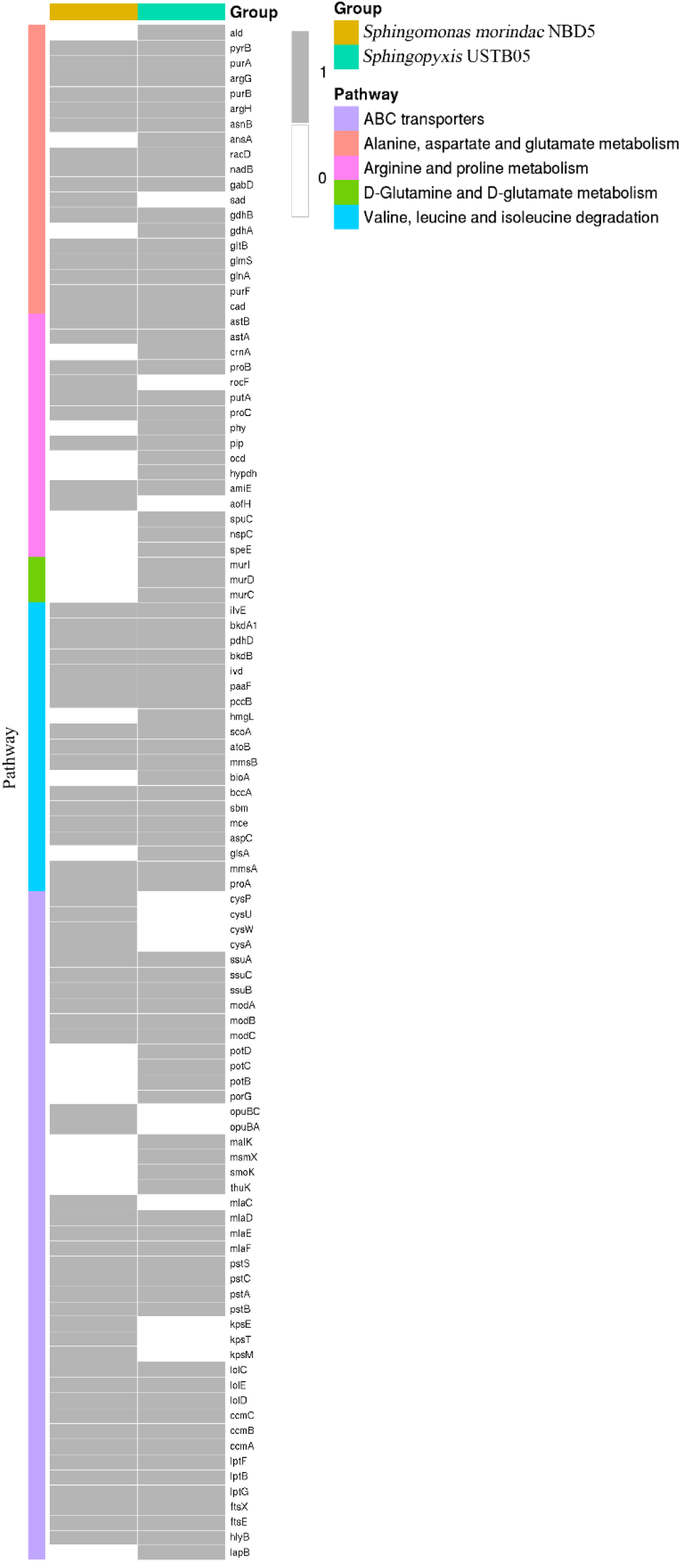
Comparison of genes related to microcystin degradation in the genomes of *Sphingomonas morindae* NBD5 and *Sphingopyxis* USTB-05. The gray box in the figure indicate that the gene exists in two strains, and the white indicate that the gene does not exist in two strains.

### Genome sequencing data comparison and proteins prediction

Based on the genome sequencing data of *Sphingomonas morindae* NBD5 and *Sphingopyxis* USTB-05, the protein sequences were predicted. The same protein sequences were combined into a cluster, and then the unique protein sequence was annotated. 1983 of protein sequences were predicted in *Sphingopyxis* USTB-05, and 1939 protein sequences were predicted in *Sphingomonas morindae* NBD5. Among them, there were a total of 1784 protein sequences showed homology, 199 unique protein sequences for *Sphingopyxis* USTB-05, and 155 unique protein sequences for *Sphingomonas morindae* NBD5 (Figure 11).

**Figure 11:**
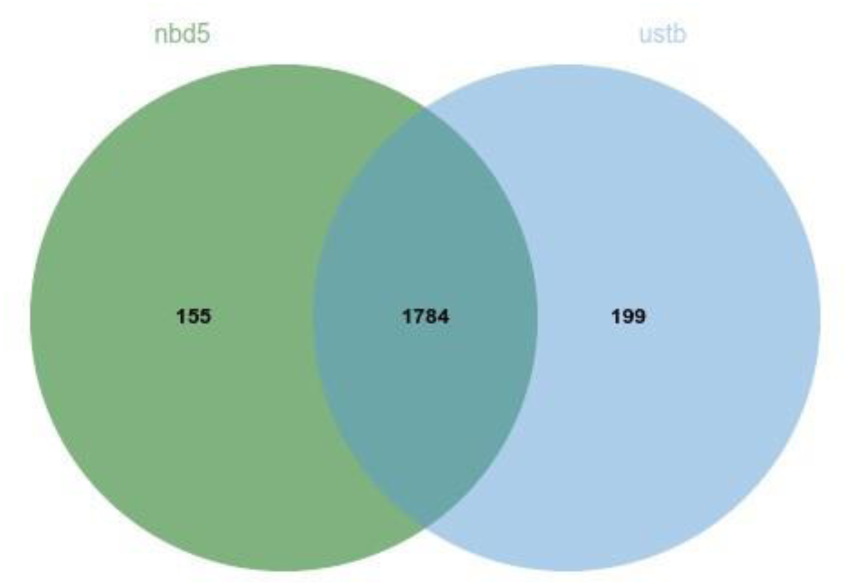
Venn diagram of comparison of genomic predicted difference proteins *Sphingomonas morindae* NBD5 and *Sphingopyxis* USTB-05.

## DISCUSSION

The two strains were found in different habitats, but they were similar or different in some functions. *Sphingomonas morindae* NBD5 had two genomes, which were relatively rare in *Sphingomonas*. The genome and plasmid prediction showed two genomes and two plasmids, indicated that *Sphingomonas morindae* NBD5 was a special strain of *Sphingomonas*. According to the description in the Berger’s Handbook of Bacteria, the GC mole percentage of *Sphingomonas* was 59-68% [7]. The GC content of the genome of *Sphingomonas morindae* NBD5 was 70%.

Genes exchanged between various species occurs frequently. The GC content of both genomes of *Sphingomonas morindae* NBD5 was 70%, and the GC content of its plasmids was 63%. The results indicated that the plasmids of strain NBD5 may have been obtained from other species during evolution. In addition, the GC content of *Sphingopyxis* USTB-05 was 64%, which was close to the GC content of plasmids of strain NBD5.

*Erwinia uredovora* was a representative of carotenoid production by prokaryotes, and its carotenoid synthesis coding genes had been studied. The carotenoid biosynthesis gene clusters with GGPP as the precursor contain 6 open reading frames (ORFs), which are *crtE*, *crtX*, *crtY*, *crtI*, *crtB* and *crtZ*. α-carotene is converted into zeaxanthin by β-carotene 3-hydroxylase, which could be derived from the *crtZ* of *Erwinia uredovora* and the *CrtR-b* of the two strains of this study (16). In addition, there was a protein-coding gene *CrtL-b* that catalyzed the mutual conversion of different types of carotene in the lutein synthesis and metabolism pathway of the two strains in this study (Figure 8).

Combining the synthetase genes were found in the terpenoid backbone biosynthesis pathway (Figure 7) and carotenoid biosynthesis pathway (Figure 8), the most probable lutein synthesis pathway was predicted. In these two strains of Sphingomonadaceae, 1-deoxy-_D_-xylulose 5-phosphate (DXP) is synthesized from pyruvate and _D_-glyceraldehyde 3-phosphate via the 2-Cmethyl-_D_-erythritol 4-phosphate (MEP) pathway. After seven steps of enzyme reactions, geranyl diphosphate (GPP) or isopentenyl pyrophosphate (IPP) is generated. Taking GGPP as the precursor, lutein is synthesized by nine steps of enzyme reactions. These reaction products are prephytoene pyrophosphate, phytoene, phytofluene, ξ-carotene, neurosporene, lycopene. Then, δ-carotene or α-carotene is reacted with zeaxanthin or α-cryptoxanthin, and hydroxylase is hydroxylated to form lutein. Among them, the metabolic process of synthesizing GGPP from GPP or IPP is lack. However, the molecular structures of zeaxanthin and lutein are only different in the position of the double bond of the left six-membered carbon ring, and they can be synthesized via the 2-Cmethyl-_D_-erythritol 4-phosphate (MEP) pathway. The synthesis pathway of zeaxanthin can provide an example. The condensation of IPP with dimethylallyl diphosphate (DMAPP) to form GPP is catalysed by geranyl diphosphate synthase (GPS). GPP is condensed with one molecule of IPP to form farnesyl diphosphate (FPP) by FPP synthase (FPS). One molecule of FPP condenses with one molecule IPP to form GGPP under the catalysis of GGPP synthase (*CrtE*) (17)(18). Last but not least, whether there were new enzymes in the metabolic process of synthesizing carotene from lycopene was worthy of further investigation.

Hepatoxin MCs degrading bacterium were found in *Arthrobacter* spp., *Brevibacterium* sp. and *Rhodococcus* sp. (19). However, the presence of biodegradable hepatoxins MCs in Sphingomonadaceae has been reported more frequently (15). Through cloning and gene library screening, the gene clusters for biodegrading MC-LR (*mlr A*, *mlr B*, *mlr C* and *mlr D)* were identified preliminarily by Bourne et al (20). It was further discovered that all or part of these four genes existed in many MCs biodegrading strains. Molecular research found that *Sphingopyxis* USTB-05 contained *USTB-05-A*, *USTB-05-B*, *USTB-05-C* genes with high homology to *mlr A*, *mlr B*, *mlr C*, respectively (21). Hashimoto et al. (22) speculated that the genes involved in the degradation of MCs were far more than these four genes. Through the KEGG database metabolic pathway annotation, the following genes may be involved in the biodegradation process of hepatotoxins MCs and NOD by *Sphingopyxis* USTB-05. These genes *ald*, *ansA* and *gdhA* in the metabolic pathways of alanine, aspartate and glutamate are involved in the biodegradation of D-alanine, D-erythro-β-methylaspartate and D-glutamate. These genes *crnA*, *phy*, *ocd*, *hypdh*, *spuC*, *nspC* and *speE* in the metabolic pathway of arginine and proline are involved in the biodegradation of L-arginine. These genes *murI*, *murD* and *murC* in the metabolic pathways of D-glutamine and D-glutamate are involved in the biodegradation of D-glutamate. These genes *hmgL*, *bioA* and *glsA* in the metabolic pathways of valine, leucine and isoleucine are involved in the biodegradation of D-isoleucine (Figure 10). *Sphingopyxis* USTB-05 has the function of biodegrading MCs, but *Sphingomonas morindae* NBD5 does not. *Sphingopyxis* USTB-05 has 199 unique protein sequences, including MCs degradation-related unique aspartate-type endopeptidase activity (GO: 0004190), metallopeptidase activity (GO: 0004181) and carboxylate hydrolase activity (GO: 0052689).

The biodegradation pathway of *Sphingopyxis* USTB-05 for MC-YR and NOD has been clarified. The first step is to convert the cyclic structure to the linear structure by the same enzyme. Adda is produced as the final product by their last enzyme reaction. However, the further biodegradation of Adda by *Sphingopyxis* USTB-05 has not been reported yet. Recently, these genes and transposable elements that may be involved in the biodegradation of phenylacetate have been observed near the *mlr* gene cluster (23). *Sphingopyxis* sp. YF1, which relies on the *mlr* biodegradation pathway, can biodegrade Adda through phenylacetic acid metabolism (24). Through the KEGG database metabolic pathway annotations, the evidence of further biodegradation of Adda is found. The biodegradation of Adda is related to some degradation genes in the metabolic pathways of styrene, toluene and xylene (Suppl Figure S1-3).

The results of *Sphingobium indicum* B90A, *Sphingobium japonicum* UT26 and *Sphingobium francense* Sp+ show that they are able to transform β- and δ-hexachlorocyclohexane (β- and δ-HCH, respectively), the most recalcitrant hexachlorocyclohexane isomers, to pentachlorocyclohexanols, but only *Sphingobium indicum* B90A can further transform the pentachlorocyclohexanol intermediate to the corresponding tetrachlorocyclohexanediols (25). The *linB* gene of *Sphingobium indicum* B90A heterologously expressed protein was incubated with γ- and β-hexachlorocyclohexane, the pentachlorocyclohexanol product was further transformed and eventually disappeared from the culture medium (25). The *linB* gene was also found in the annotation of chlorocyclohexane and chlorobenzene metabolism of *Sphingopyxis* USTB05 (Suppl Figure S4). In addition, *Sphingopyxis* USTB05 also found many genes related to the degradation of environmental pollutants, such as: dioxins, toluene, xylene, chlorocyclohexane and chlorobenzene, chloroalkane and chloroalkene, styrene, naphthalene and other degradation genes (Suppl Figure S5-7). Although the functions of these genes in the *Sphingopyxis* USTB05 genome still need to be verified, it still shows that *Sphingopyxis* USTB05 has great potential as an environmental pollutant degradation bacteria.

## MATERIALS AND METHODS

### Bacterial strains and culturing conditions

*Sphingopyxis* sp. USTB-05 was isolated and identified from the sediment of Dianchi Lake in China (5). *Sphingomonas morindae* NBD5 was isolated and identified from Noni (*Morinda citrifolia* L.) branch (4). They were grown in LB media at 30 °C.

### DNA extraction, identification and sequencing

Glycerol stocks of the original strains were initially used as inoculum for regrowth on the original solid isolation media at 30 °C. Single colonies were picked and cultured in LB media. Genomic DNA was extracted using the Bacterial Genomic DNA Kit (CoWin Biosciences, China) according to the manufacturer’s instructions. DNA quality and integrity were checked by NanoDrop Nucleic Acid Quantification (Thermo Fisher Scientific, USA) and gel electrophoresis.

To confirm the identity of the strains, 16S ribosomal RNA (rRNA) gene amplicons were generated by PCR using primers 27F (5′-AGAGTTTGGATCMTGGCTCAG-3′) and 1492R (5′-GGTTACCTTGTTACGA CTT-3′). The PCR reaction mixture contained 25 μL PCR Mix, 2 μL primer 27F (10 mM), 2 μL primer 1492R (10 mM) and 1μL of the extracted DNA. Nuclease-free water was added to reach a total reaction volume of 50 μL. The following conditions were used for the bacterial 16S rRNA gene amplification: initial denaturation at 94 °C for 5 min followed by 30 cycles of denaturation at 94 °C for 40 s, annealing at 55 °C for 40 s, elongation at 72 °C for 45 s and a final extension step at 72 °C for 10 min. PCR products were purified using 1% gel electrophoresis. The purified PCR products were sent for DYY-8C DNA sequencer sequencing. The sequencing results was put in EZ biocloud Alignment to determine the homology relationship with the known sequence.

The genomic DNA of two strains was constructed by nanopore single molecule sequencing library according to the standard protocol provided by Oxford Nanopore Technologies (ONT). Large fragments of DNA were recovered by using BluePippin (Sage Science, USA) automatic nucleic acid recovery system (26). DNA damage repair and end repair, magnetic beads purification and linker connection were processed by using the official SQK-LSK 109 ligation kit (Oxford Nanopore, UK). The Qubit library quantification were transferred to computer sequencing. A second-generation sequencing library for the genomic DNA and plasmid DNA of the two strains was constructed. Genome sequencing of strains USTB-05 and NBD5 was performed using the Illumina MiSeq platform (paired end, 2 × 300 bp reads) (27).

### Genome assembly and quality control

For genome assembly, the subreads used second-generation sequencing technology were first filtered with fastp software to obtain high-quality reads. The sequences containing the linker fragment were deleted. The sequences with mass value Q <25 and low mass base number length more than half of the sequence length were deleted.

For nanopore data, the original fastq format was obtained by base calling fast 5 file through Albacore software in MinKNOW software package. In order to obtain more accurate assembly of results, it was necessary to filter these impurities to obtain reliable subreads, including filtering out Polymerase Reads with a length less than 1000 bp.

### Genome annotation and comparative genomic analysis

The online NMPDR-rust server was used to predict the gene and coding sequence (CDs) region of the assembly sequence. The predicted protein sequences were searched against KEGG (Kyoto Encyclopedia of Genes and Genomes), COG (Cluster of Orthologous Groups), GO (Gene ontology) to predict gene functions and metabolic information through Blastall (28). Circos software was used to integrate the COG annotation results, methylation results, RNA annotation results, GC content, and GC-skew to map the entire genome of the bacterial strain. In addition, CRISPRFinder software was used to predict the clustered regularly interspaced short palindromic repeats (CRISPR) structure of the genome (29). The coding sequences of the two genomes were aligned using MUMmer and analyzed in conjunction with the results of the genome annotation (30).

### Drawing tool

The heat map was generated by using the online software Hiplot (https://hiplot.com.cn). Adobe Illustrator CS6 was used to generate other figures.

## ABBREVIATIONS

AMD: age-related macular degeneration
MCs: microcystins
NOD: nodularin
WHO: world health organization
DD: dibenzo-p-dioxin
LPS: lipopolysaccharide
CDSs: coding sequence
CRISPR: clustered regularly interspaced short palindromic repeats
GO: gene ontology
COG: cluster of orthologous groups
KEGG: kyoto encyclopedia of genes and genomes
VFDB: virulence factors database
CARD: comprehensive antibiotic resistance database
ORF: open reading frame
NMEP: non-mevalonate pathway
GPP: geranyl pyrophosphate
IPP: isoprene pyrophosphate
GGPP: geranyl geranyl pyrophosphate

## ACKNOWLEDGMENTS

This work was supported by the National Natural Science Foundation of China (21677011) and the Fundamental Research Funds for the Central Universities (FRF-TP-20-044A2; FRF-BR-19-003B; FRF-TP-18-012A1; FRF-TP-17-009A2).

## Supplementary Tables and Figures

**Suppl Table S1.**
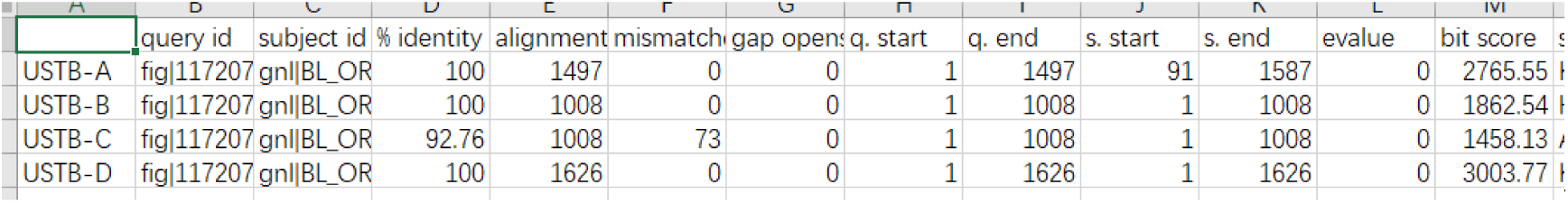
The sequences of *USTB-A*, *USTB-B*, *USTB-C*, and *USTB-D* in the *Sphingopyxis* USTB-05 genome are similar to *mlr A*, *mlr B*, *mlr C* and *mlr D*, respectively.

**Suppl Figure S1.**
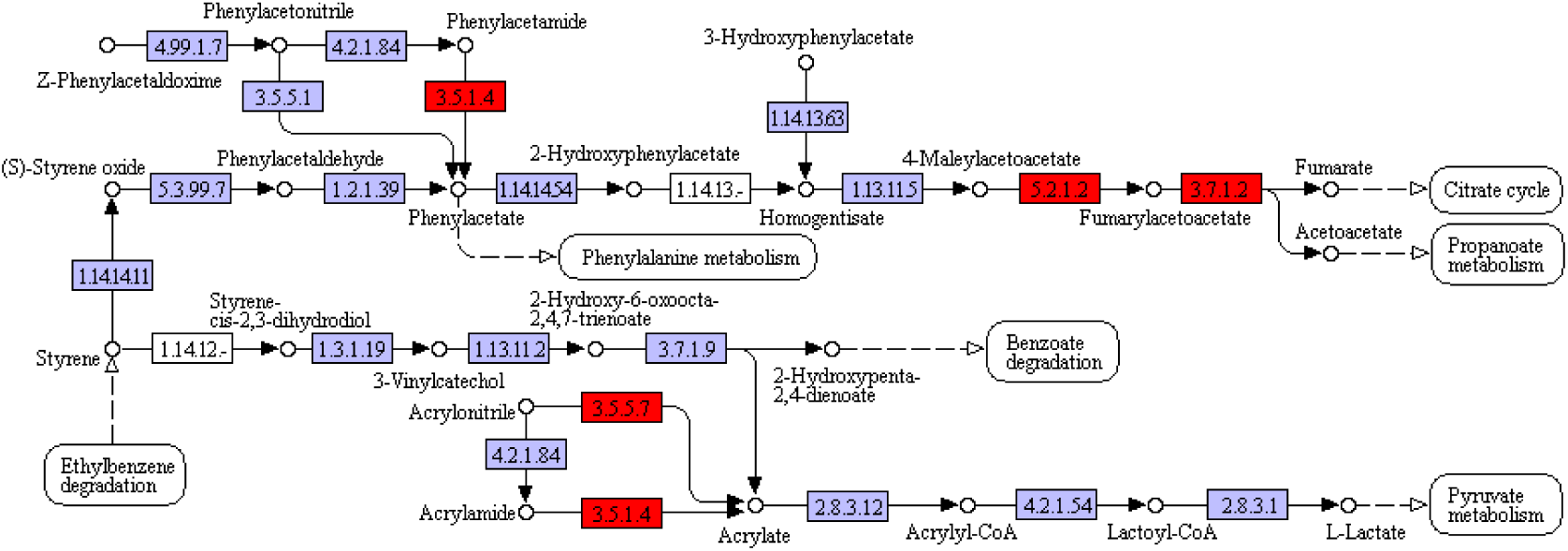
The metabolic pathway of styrene degradation of *Sphingopyxis* USTB-05.

**Suppl Figure S2.**
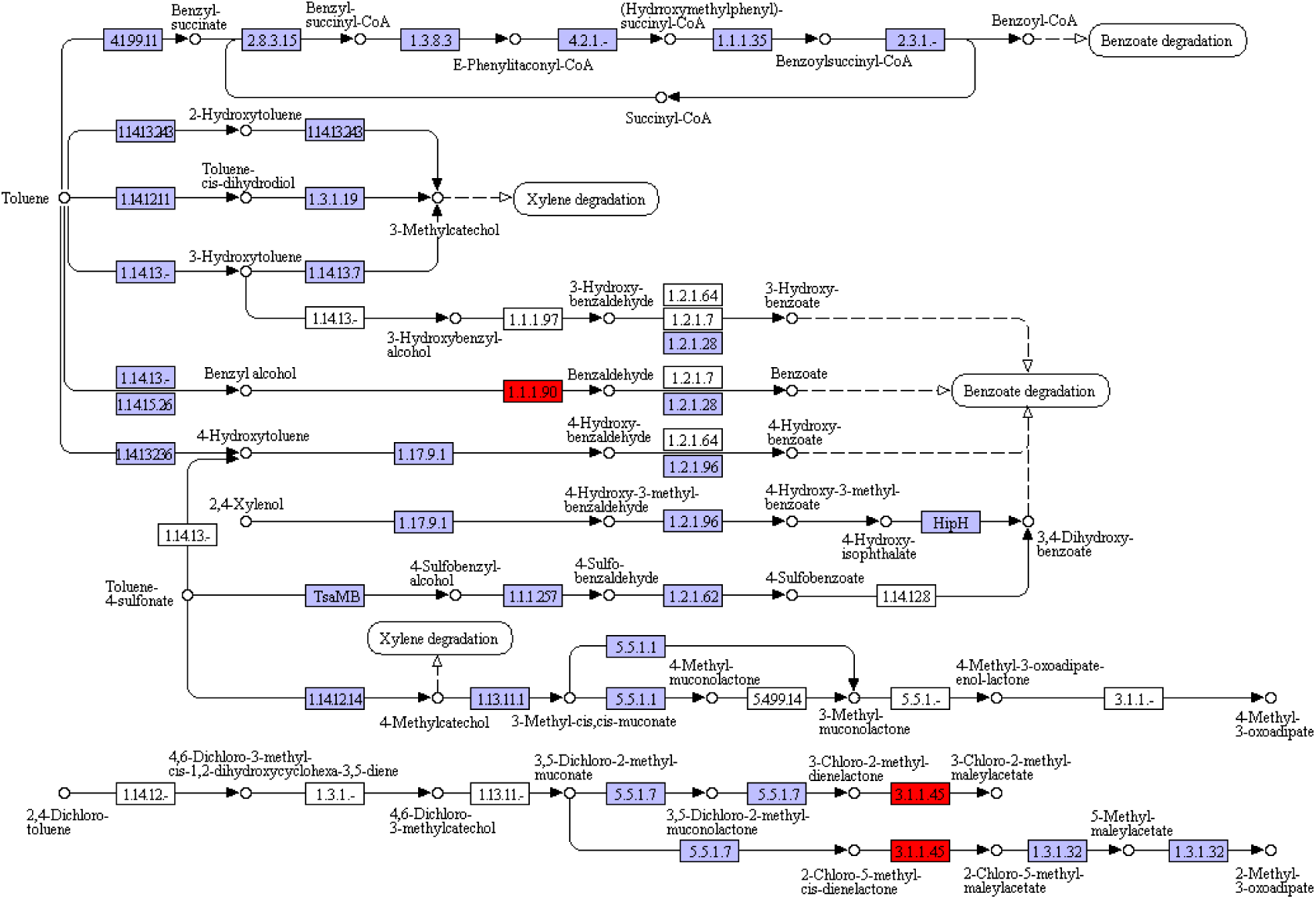
The metabolic pathway of toluene degradation of *Sphingopyxis* USTB-05.

**Suppl Figure S3.**
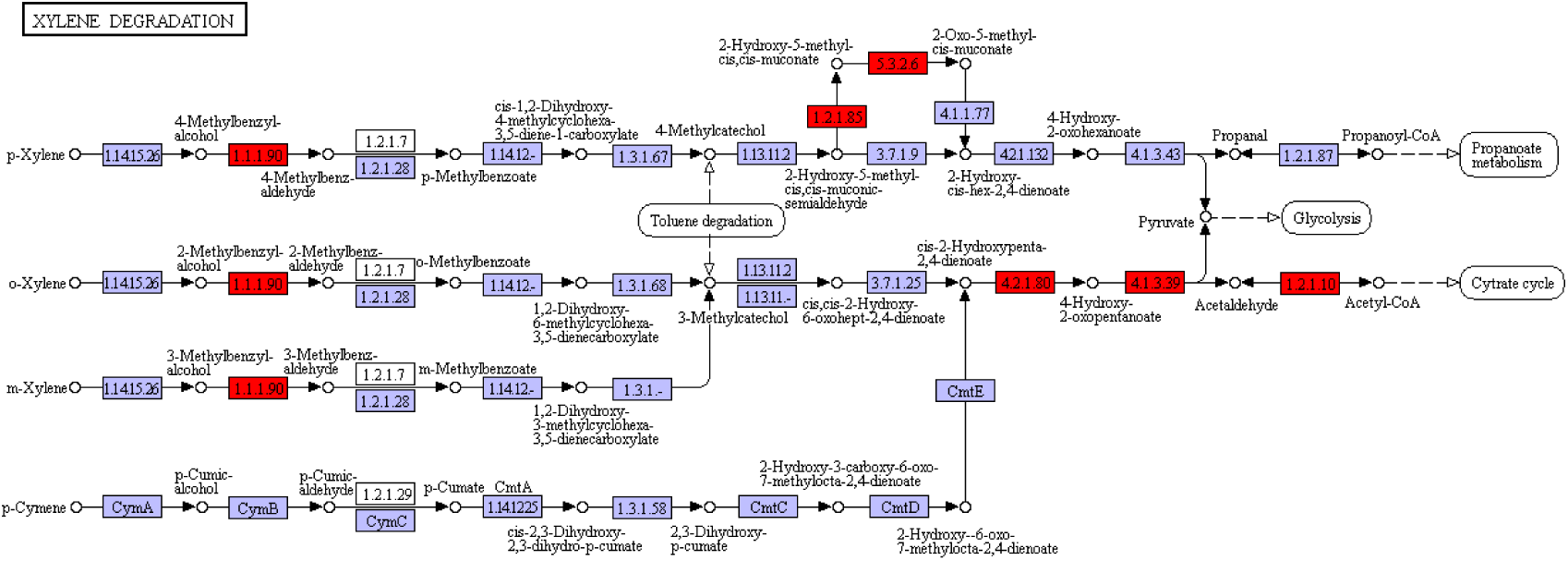
The metabolic pathway of xylene degradation of *Sphingopyxis* USTB-05.

**Suppl Figure S4.**
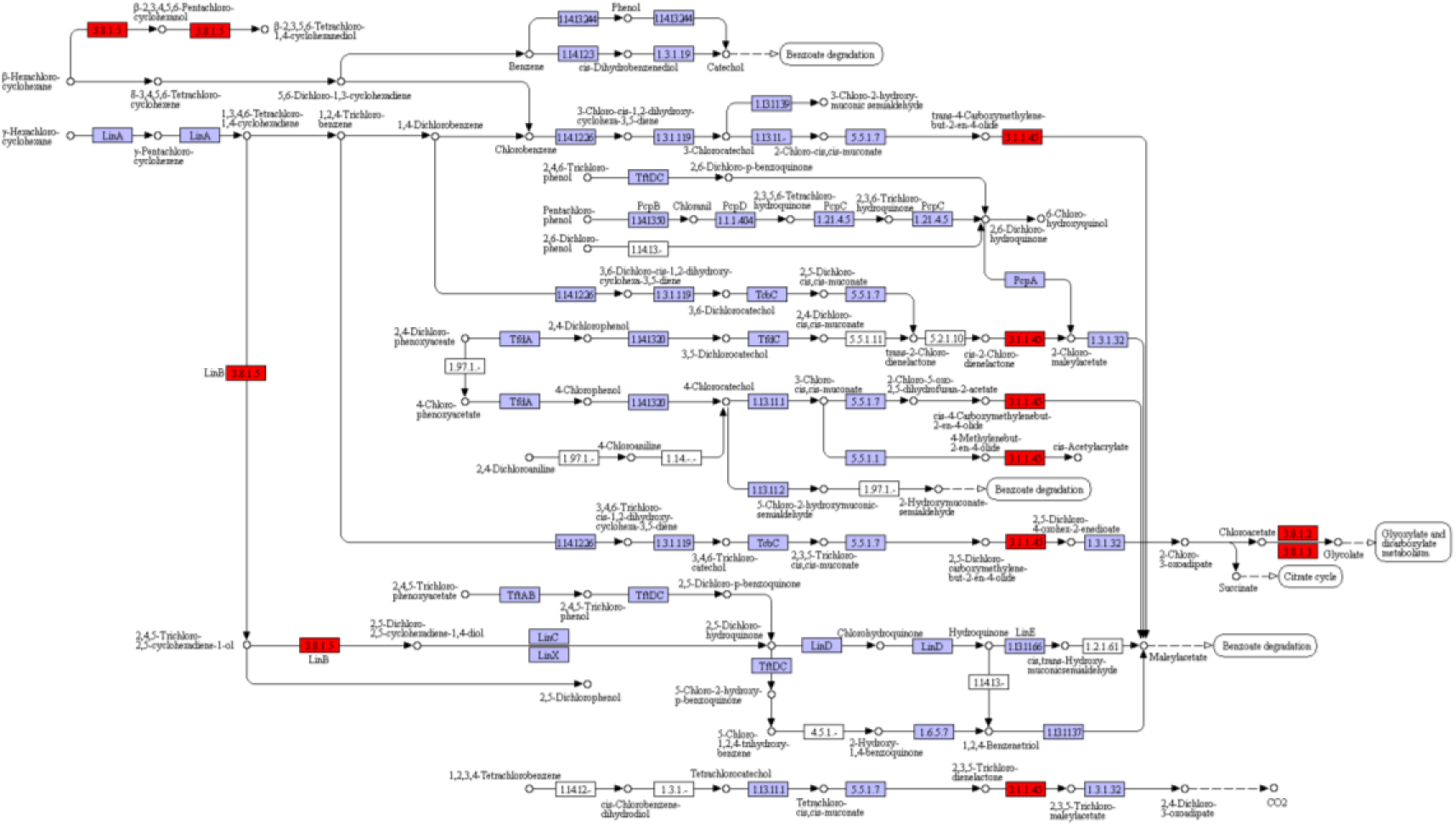
The metabolic pathway of chlorocyclohexane and chlorobenzene degradation of *Sphingopyxis* USTB-05.

**Suppl Figure S5.**
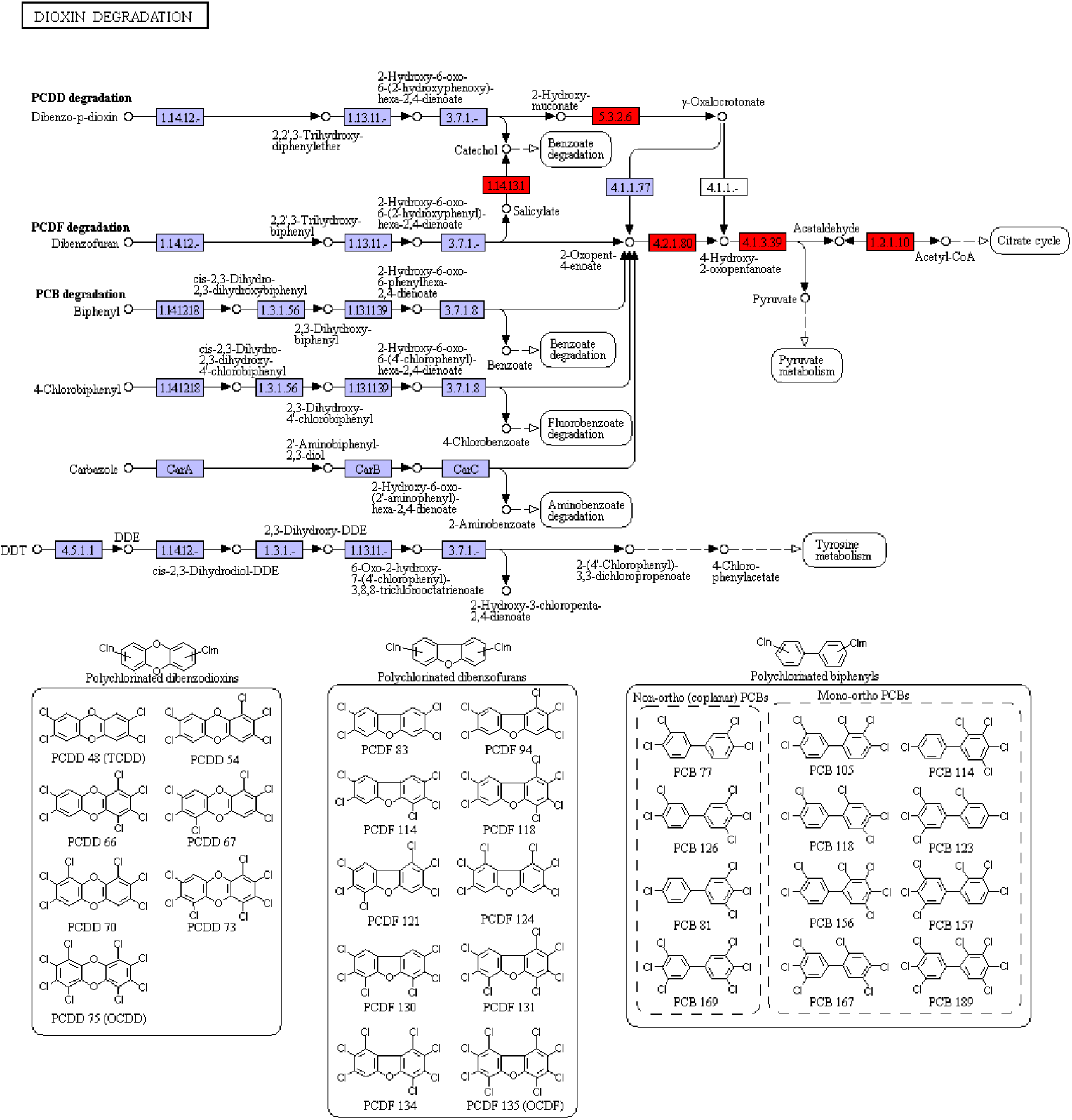
The metabolic pathway of dioxin degradation of *Sphingopyxis* USTB-05.

**Suppl Figure S6.**
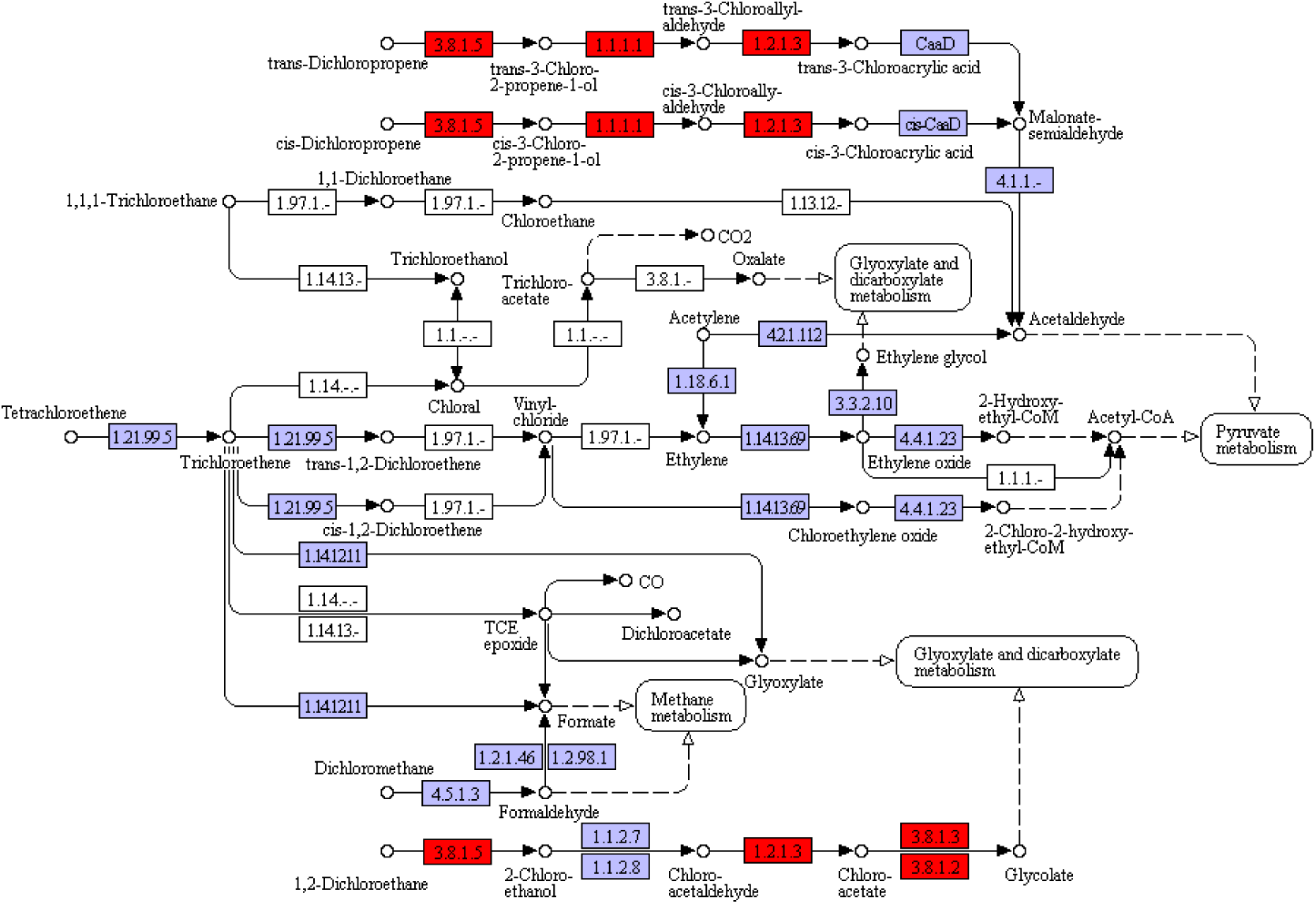
The metabolic pathway of chloroalkane and chloroalkene degradation of *Sphingopyxis* USTB-05.

**Suppl Figure S7.**
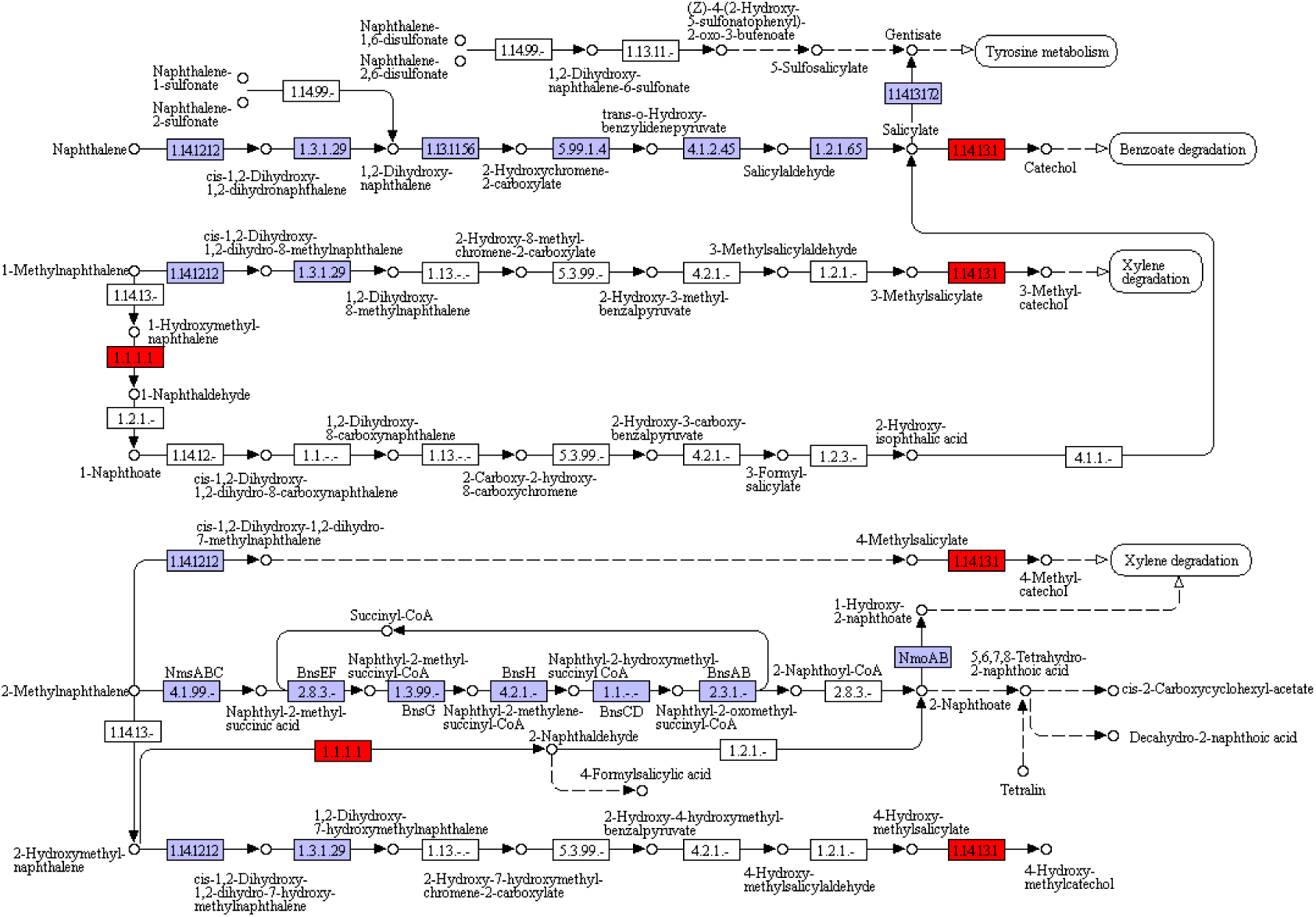
The metabolic pathway of naphthalene degradation of *Sphingopyxis* USTB-05.

